# Genetic Determinants of Heart Failure Susceptibility and Response in the Collaborative Cross Mouse Population

**DOI:** 10.1101/2025.10.07.681046

**Authors:** Todd H. Kimball, Anh N. Luu, Brian Gural, Caitlin Lahue, Abigail Hockett, Sriram Ravindran, Amira Ali, Aryan Dalal, Sam Ardery, Emily L. Sipko, Logan G. Kirkland, Mansi Goyal, Brian C. Jensen, Rebecca B. Berlow, Christoph D. Rau

## Abstract

Genetic variation and lived experiences shape how our hearts respond to chronic stress. The specific genetic mechanisms which underly cardiac remodeling, however, are still unclear, due in part to the challenge of accounting for environmental effects in human population studies. To overcome this challenge, we used the Collaborative Cross (CC) mouse population to investigate heritable susceptibility to cardiovascular stress by chronic β-adrenergic receptor stimulation.

Across 8 founder and 63 CC lines, we measured cardiac structure and function, organ weights, cell and tissue morphology, and left ventricular gene expression. Genome-wide scans detected 49 genome-wide significant loci, collapsing to 20 unique intervals (nine significant for multiple traits and eleven trait-specific), averaging 12.83 Mb in size.

To identify high-confidence candidate genes from these loci, we augmented our trait mapping with associations between loci and gene expression, isoproterenol-dependent transcriptional changes, coding variants drawn from sequencing data, tractability in our *in vitro* rat cardiomyocyte model, and previously reported protein functions and mouse or human phenotypes. This approach recovered both known regulators, such as *Hey2*, and new candidates.

Functional tests in *in vitro* models highlight three candidate genes that modulate hypertrophic growth: *Abcb10*, *Mrps5* and *Lmod3*. *Abcb10* knockdown increased cell size at baseline and further with isoproterenol, consistent with loss of a mitochondrial stress-buffering role. *Mrps5* knockdown blunted stress-induced hypertrophy. Paradoxical upregulation of *Lmod3* after siRNA transfection (validated at the protein level) also attenuated hypertrophy, consistent with reinforcement of actin-assembly control under catecholamine stress.

Together, these results reveal heritable pathways of β-adrenergic remodeling in mice and provide an interpretable, translational, and stepwise framework to prioritize candidate genes within broad loci for mechanistic studies of heart failure.

## Introduction

At some point in our lives, we will be asked if heart disease runs in the family. For clinicians, family history is among the most important factors^1^ in understanding if their patient is at risk of joining the 30 million people worldwide affected by heart failure^2^, a chronic condition in which the heart is unable to pump sufficient blood to meet the body’s needs. In 2020, an estimated 6.7 million Americans possessed a heart disease diagnosis and by 2030, the forecasted number will rise to 8.5 million^3,4^. The crucial Framingham Heart Study illuminated risk factors that predispose individuals to developing heart failure, including age, smoking and obesity^4^, while other epidemiological studies have shown the socio-economical effect on specific cardiac etiologies, including ischemic, hypertensive, and valvular heart disease^1^. Underlying these environmental factors is the genetic heritability of heart failure (estimated in human populations at ∼26%^5^), which acts to amplify or diminish risk factor significance or the etiological basis of diagnosis. Even as the tools to identify genetic contributors to heart disease have improved: easier access to data sets, increased levels of population diversity^2^, and the successful algorithmic identification of rare, less penetrative causal mutations^6^, the sheer number of contributing factors to an individual diagnosis in humans decreases the chances of discovering and linking specific allelic variants with heart failure risk.

Murine genetic reference populations (GRPs) are purpose designed for studying multi-factor traits like heart failure. They enable direct access to diseased tissues, complete environmental control, and, in many cases, treatment-control comparisons between genetically identical replicates. The Collaborative Cross (CC), a widely used GRP, is a large panel of incipient inbred lines derived from eight founder strains that allows for phenotypic characterization in a controlled, genetically diverse and repeatable manner^7^. The CC has been utilized to genetically map genomic regions and genes linked to diseases and phenotypes including host-pathogen interactions^8^, immunological conditions^9^, and ozone exposure^10^. Recently, two reports have used the CC to study the role of genetics in myocardial infarction^11^ and baseline variation in echocardiographic measurements^12^. There is a pressing need to expand on these studies by examining the effects of chronic stress on the heart and its progression towards heart failure, a gap unexplored by either of these prior two CC cohorts.

Heart disease is a progressive disorder^13^ governed by multiple neurohormonal signaling mechanisms, including the sympathetic nervous system (SNS)^14^, renin-angiotensin axis^15^, and endothelin axis^16^, all which act to modulate cardiac workload and output to maintain organismal homeostasis. The SNS releases the catecholamines epinephrine and norepinephrine which act on β-adrenergic receptors on cardiomyocytes to increase contractile rate and force^17^. This compensatory signaling remodels the cardiac tissue, increasing left ventricular wall thickness to maintain cardiac output, however prolonged adrenergic activation becomes maladaptive as initial hypertrophic growth may lead to obstruction and or eventual dilation of the left ventricle^18^, fibrotic deposition^19^, and inflammation^20^. These responses are underlined by transcriptomic responses that further propagate pathophysiological mechanisms, including metabolic dysfunction and sarcomeric protein reorganization. Treatment of the CC lines with the β-adrenergic agonist isoproterenol (ISO) induces a spectrum of phenotypic and transcriptomic outcomes and mapping these traits to genetic variations allows us to identify genetic loci associated with cardiac disease^21,22^.

In this study, we subject mice from 63 strains of the CC as well as the 8 CC founder lines to ISO treatment to induce cardiac hypertrophy and, in some cases, heart failure. We then utilize state-of-the-art GWAS approaches tailored to the CC population to identify loci which drive cardiac phenotypes with or without ISO stimulation. By doing so, we utilize the CC’s significant genetic diversity and improved mapping capabilities to augment previous discoveries and guide attention to novel candidates for therapeutic development.

## Methods

### Animal Husbandry

All animal experiments were conducted under protocols established and approved by the University of North Carolina at Chapel Hill Institutional Animal Care and Use Committee. (CC portion: 22-220.0, NRVM portion: 22-217.0) All surgeries were performed under isoflurane anesthesia, and every effort was made to minimize suffering. Human endpoints included loss of weight, lethargy, hunched posture, and other standard measures. Animals were kept in hot-washed cages within the vivarium, monitored daily by vivarium staff, and provided standard chow and enrichment. We adhered to the ARRIVE guidelines for this study.

### Animals

#### Collaborative Cross Mice

Mice were procured from the UNC Systems Genomics Core at an age of 6 weeks and allowed to acclimatize to our vivarium for three weeks preceding pump implantation. See **Supp. Table 1** for the names of all mouse strains used in this study.

#### Rats

Wistar rats were procured from Charles River and bred for NRVM isolation from pups (detailed below).

### Chronic β-adrenergic Stimulation *in vivo*

Mice were weighed and ALZET Micro-osmotic Pumps (Model 1004) were loaded with 30 mg/kg body weight/day of ISO (or saline for sham mice). Mice were anesthetized and the pump was implanted subcutaneously by the UNC McAllister Heart Institute Cardiovascular Physiology and Phenotyping Core.

### Echocardiography and Phenotypic Measurements

A Vevo-F2 ultrasonic imaging system (VisualSonics) was used to perform echocardiography on anesthetized mice (1-2% isoflurane in 98% O2, 2% CO2) and their heart rate was monitored and isoflurane titrated to achieve imaging between 450–550 bpm. Measurements were taken at basal timepoint before pump implantation and 21 days following. Images of the left ventricular short axis were captured in B-mode and M-mode to measure cardiac physiology. Measurements were performed using the Vevo-Lab software suite for left ventricular wall thickness and chamber size to calculate ejection fraction, fractional shortening, and other cardiac parameters. See **Supp. Table 2** for details on all echocardiographic and other phenotypic readouts.

### Tissue Collection and Preparation

Mice were euthanized 24 hours after the 21-day echocardiogram. Their body weight was recorded, the heart removed and weighed, then dissected into the 4 chambers to be subsequently individually weighed. The lungs, liver, and adrenal gland were also removed and weighed. The left ventricle was further dissected by two transverse cuts in the midsection, with the middle section reserved for histological studies. All other tissue samples were flash frozen in liquid nitrogen followed by storage at -80° C. The histological samples were placed in formalin for at least overnight to preserve cardiac ultrastructure. Samples were embedded in paraffin and stained with Masson’s Trichrome and H&E by the UNC Pathology Services Core Facility. Additional slides were prepared by cutting 5 µm slices with a microtome and dried at 37° C overnight. To assess cardiac fibrosis, tissue sections were stained with Sirius Red/Fast Green Collagen Staining Kit (Chondrex Cat #9046) and whole tissue sections were imaged with the ECHO Revolve RVL-100-M microscope with polarizing film sheets (Amazon Item model 93493) to produce polarized images. Assessment of fibrotic deposition percentage to whole tissue was calculated with the FibroSoft software package^23^. Cardiomyocyte cross-sectional area was calculated by staining tissue sections with Wheat Germ Agglutinin (WGA) per the manufacturer protocol and imaging at 20x with DAPI co-stain. A total of 10 fields and 100 cells were traced via ImageJ and average cross-sectional areas calculated for each sample.

### RNA Isolation and Library Prep

RNA was isolated from left ventricular tissue samples with QIAGEN RNA-Mini Kit (Cat: 74106). RNA purity was assessed with a nano-drop to achieve quality 260/230 ratio > 1.6. RNA samples were additionally assessed via TapeStation for RIN > 7.

Five RNA libraries representing 440 total samples were prepared for bulk sequencing using the Genomics Mercurius Full Length mRNA BRB-seq kit (Cat: 11013 and 10813) per manufacturer guidelines. Each library was prepared on a 96-well plate, where 50 ng RNA was combined with well-specific barcodes and pooled together for cDNA synthesis. Samples were assessed for quantity, indexing barcodes were added to ensure 5 unique libraries, and each amplified through 12 cycles of PCR. Final libraries were assessed for RNA integrity by Tapestation.

### Mouse Genotypes and Sequences

CC Genomes were obtained from the CC Mouse Genotypes Resource (https://www.csbio.unc.edu/CCstatus/CCGenomes/)^24^ and converted into a HAPPY-style genome cache for downstream analyses^25^. Sequences of the Collaborative Cross founders were obtained from the Wellcome Trust Mouse Genomes Resource (https://www.mousegenomes.org/)^26^ and filtered for SNPs predicted to cause a non-synonymous mutation in a protein coding gene.

### RNA Sequencing + Transcriptomic Analysis

Sequencing was conducted by the Sequencing and Genomic Technologies Shared Resource in the Duke University School of Medicine. Five sample libraries were sequenced by paired end sequencing on a single Novaseq X 25 Gb chip.

Raw reads were returned from sequencing in fastq format and read quality was checked with FastQC^27^ and summarized with MultiQC^28^. Following the manufacturer recommendations, fastq files originating from the same library plate were then concatenated together to facilitate identification of sample-specific barcodes within pooled libraries (demultiplex). Next, we used STAR v2.7.9a to generate an index from the Mus musculus primary assembly genome (GRCm39) and its corresponding gene transfer format (GTF) file (Ensembl release 110).

Reads from each of the five libraries were then aligned to the indexed genome leveraging STAR’s single-cell RNA-seq alignment mode (soloType CB_UMI_Simple). This process demultiplexed reads based on a 14 bp sample barcode and a 14 bp unique molecular identifier (UMI). Reads mapping to more than one genomic location were removed. STAR was also used to quantify reads per gene, producing a raw gene-cell counts matrix for each library.

The five resulting count matrices were loaded into R and merged into a single data frame. Sample metadata was then joined to the count data. We used the biomaRt package^29,30^ to convert Ensembl gene IDs to MGI gene symbols. In cases where multiple Ensembl IDs mapped to a single gene symbol, counts were summed to the gene-level.

To manage potential batch effects, sequencing libraries were prepared such that they contained technical replicates of a subset of samples which were used during normalization with ComBat-seq^31,32^ with the smaller library removed after normalization to counts per million (CPM). Only genes with an average of at least 1 CPM across all samples are analyzed further. For each treatment group, gene expression underwent sex-aware variance stabilizing transformation (VST) using the DESeq2 R package^33^.

#### Identification of differentially expressed genes (DE)

Gene expression was modeled as the sum of effects from treatment and sex using the DESeq2 R package^33^.

### Quantitative Trait Loci Mapping

To account for the multi-parental population structure and unequal founder contributions of the Collaborative Cross (CC) mice, we performed quantitative trait loci (QTL) mapping using the R package miQTL^34^. This method models phenotypes based on the mosaic of founder haplotypes using a regression-on-probabilities (ROP) approach, which simultaneously estimates the effect of all eight founder haplotypes along with a general estimate of a locus’ association with the trait^34–36^.

Prior to analysis, all phenotypic traits were normalized to better approximate a normal distribution and meet the assumptions of our statistical model. We applied a Box-Cox power transformation, with the optimal lambda calculated separately for the control and isoproterenol-treated groups for each trait using the ‘boxcox’ function from the MASS R package^37^.

We conducted genome-wide scans for each normalized trait within each treatment group by fitting the following linear mixed model with miqtl::scan.h2lmm:

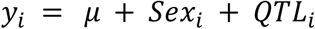

where *y*_*i*_ is the normalized trait measure for mouse *i*, *μ* is the intercept, *Sex* is the biological sex as determined by external morphology, and *OTL* is the effect of a given locus on the trait.

This model included sex as an additive covariate and incorporated a kinship matrix estimated from genome-wide haplotype data to account for the genetic relatedness between strains. Genome-wide significance thresholds were established for each trait-treatment pair via permutation testing. A null distribution was generated by performing 100 scans on randomly shuffled phenotype data, and the 90th percentile of the maximum LOD scores from these permutations was set as the significance threshold. The boundaries of significant QTL were defined by a 1.5-LOD drop from the peak marker^38^.

### eQTL Mapping

To identify genes whose expression levels were associated with the haplotypes that were also associated with phenotypic traits, we performed expression QTL (eQTL) mapping using miQTL once more, matching the design of our trait QTL mapping described above. For each gene within a significant trait locus, from treatment and control groups separately, we fit the model Gene expression ∼ 1 + Sex. Chromosome-wide significance thresholds were determined through permutation testing (n=100 permutations per gene), with the top 90% score used as the threshold LOD score. We defined a cis-eQTL as a significant eQTL where the gene resides within its own eQTL locus, and this eQTL locus, in turn, overlaps with the primary trait QTL. Genes with such colocalized signals were prioritized for further examination.

### Selection of Gene Targets

Following the scans, we identified significant QTL loci and annotated the genes within these regions using the biomaRt R package. We further characterized these genes by querying MouseMine for associated phenotypes and assessing their potential pathogenicity. All analysis steps, from the initial QTL scan to the final gene annotation, were managed and executed using a Snakemake workflow available below.

### Cell Culture Work in Neonatal Rat Ventricular Myocytes

Neonatal rat ventricular myocytes (NRVMs) were isolated from Wistar rat pups (P1-P4) using enzymatic digestion via the Worthington Neonatal Cardiomyocyte Isolation System, which uses a combination of trypsin and collagenase to release cells from minced cardiac tissue. We then performed myocyte purification by Percoll density gradient (Cytiva) where two Percoll layers (density of 1.060 g/mL and 1.082 g/mL) were used in combination with centrifugation to physically separate NRVMs from other cell types. Cells were then plated at 36,000-43,000 cells/cm^2^. After 24 hours, cells were transfected with the siRNA of our genes of interest (or scrambled sequence as control) via Lipofectamine RNAiMAX (Thermo Fisher) at a concentration of 9 pmol per 150,000 cells, then treated with 15 μM of ISO for 48 hours before formaldehyde fixation and imaging. Images were processed with Cellpose and Fiji to quantify relative cell areas^39,40^. To account for variation in raw cell size measurements between batches of rat pups, for each experimental batch, each individual cell size reading was normalized to the average cell size of the control scramble-treated group. Data for each gene underwent two-way ANOVA test followed by post hoc Tukey HSD tests.

To determine differential gene expression of candidate genes in response to ISO, NRVMs without gene knockdown were collected for RNA content post-ISO treatment. RNA was extracted (QIAGEN RNeasy Mini Kit) for downstream cDNA synthesis (Applied Biosystems™ High-Capacity cDNA Reverse Transcription Kit) and RT-qPCR (Applied Biosystems™ PowerUp™ SYBR™ Green Master Mix).

### Cell Culture Gene Knockdown Validation Assays

To validate gene knockdown, we use the H9C2 cell line (embryonic rat heart myoblasts). Cell pellets were collected 72 hours after siRNA-mediated gene knockdown, representing the original end time point in NRVM cell culture experiments. RNA extraction and RT-qPCR were conducted in a similar manner as described above. Western blot was also performed to validate change in protein levels for the *Lmod3* candidate gene that showed paradoxical mRNA upregulation after siRNA transfection. For more in detailed *in vitro* methodology, see ***Supplemental – Methods and Materials***. List of primers, siRNA, antibodies and reagent can be found in **Supp. Table 4-6**.

## Results

### Isoproterenol Induced Heterogeneous Response in the Collaborative Cross

Collaborative Cross mice, consisting of 8 founder strains and 63 derived strains (3 male and 3 female mice per strain, 444 mice in total) (**Figure 1A**), were subjected to a baseline echocardiogram (ECHO) to identify strain-specific differences in cardiac parameters. A summary of basal echocardiography phenotypes can be found in **Table 1**. Basal ejection fraction (**Figure 1C**) of CC mice demonstrated strain-specific variability, ranging from CC03 (avg = 88.64) to the founder strain C57BL/6J (avg = 46.87). Basal measurements of cardiac phenotypes, taken under anesthesia, displayed significant inter-strain variability in all measured ECHO parameters (p < 0.05). Significant sex-specific effects were observed for several echocardiographic traits for heart size at the basal timepoint, including left ventricular mass (males 16% heavier, p = 1.3⨯10^-8^), left ventricular end volume at diastole (males 16% higher, p = 9.28⨯10^-7^) and systole (males 18% higher, p = 3.0⨯10^-4^), and left ventricular internal diameter at diastole (males 6% higher, p = 2.9⨯10^-6^), but not for heart rate or ejection fraction.

**Figure 1.**
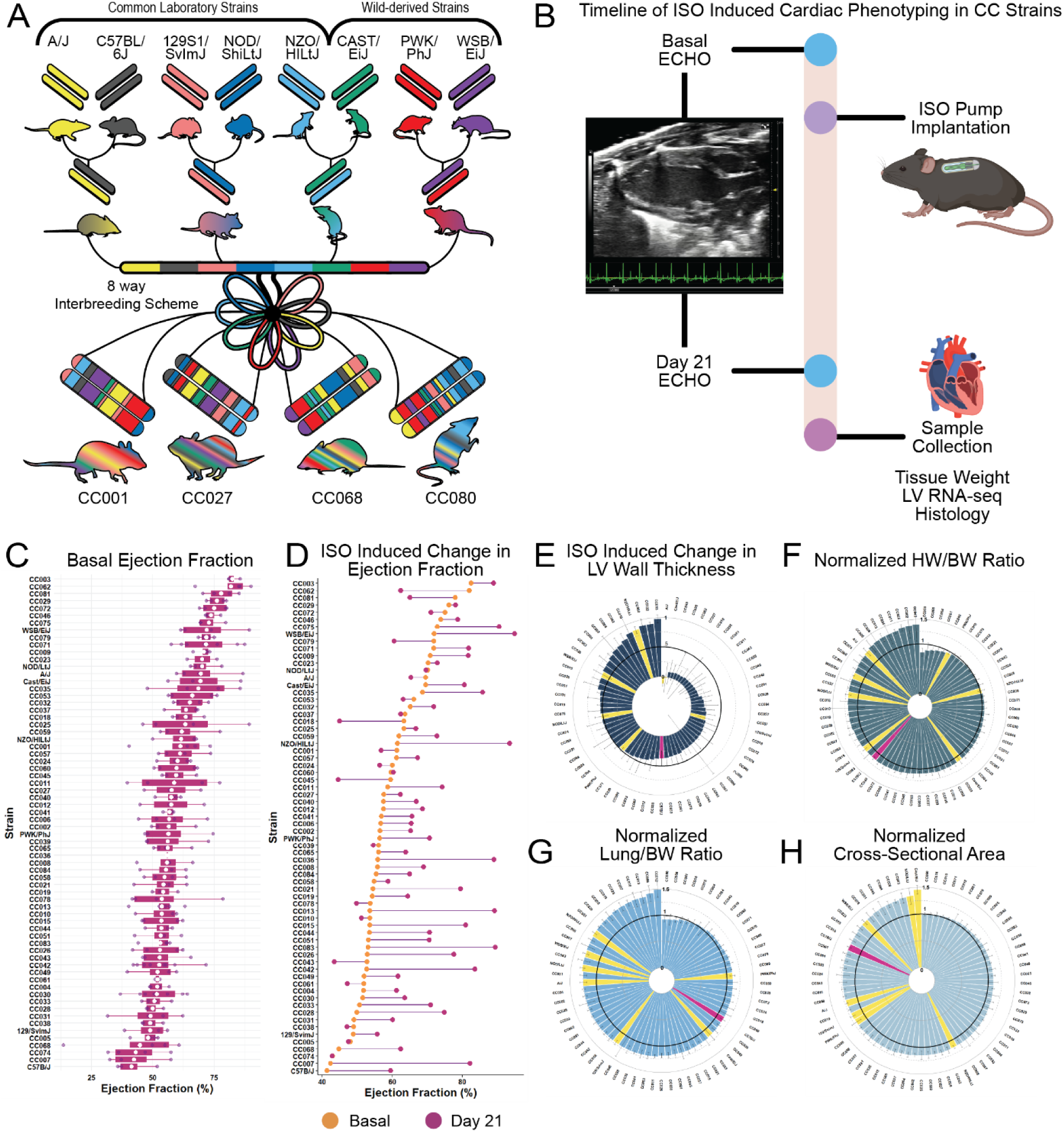
Phenotypic Characterization of Collaborative Cross Mice. **A.** Collaborative Cross strain generation schematic. Eight founder strains consisting of common laboratory strains and wild-derived strains were interbred to produce the 63 incipient inbred strains used in this study. **B.** ISO treatment and experimental timeline. Mice underwent echocardiography at basal timepoint, ISO pumps were implanted for 21 days with follow-up echocardiography. Samples were collected 24 hours after last ECHO. **C.** Average baseline ejection fraction of CC strains, white is the strain mean. **D.** ISO induced change in ejection fraction, average basal in orange, average 21 day post-ISO in purple. **E-H.** Circle plots of average phenotypic measurements, yellow and red bars are founder strains, with red being C57BL/6J.

**Table 1:**
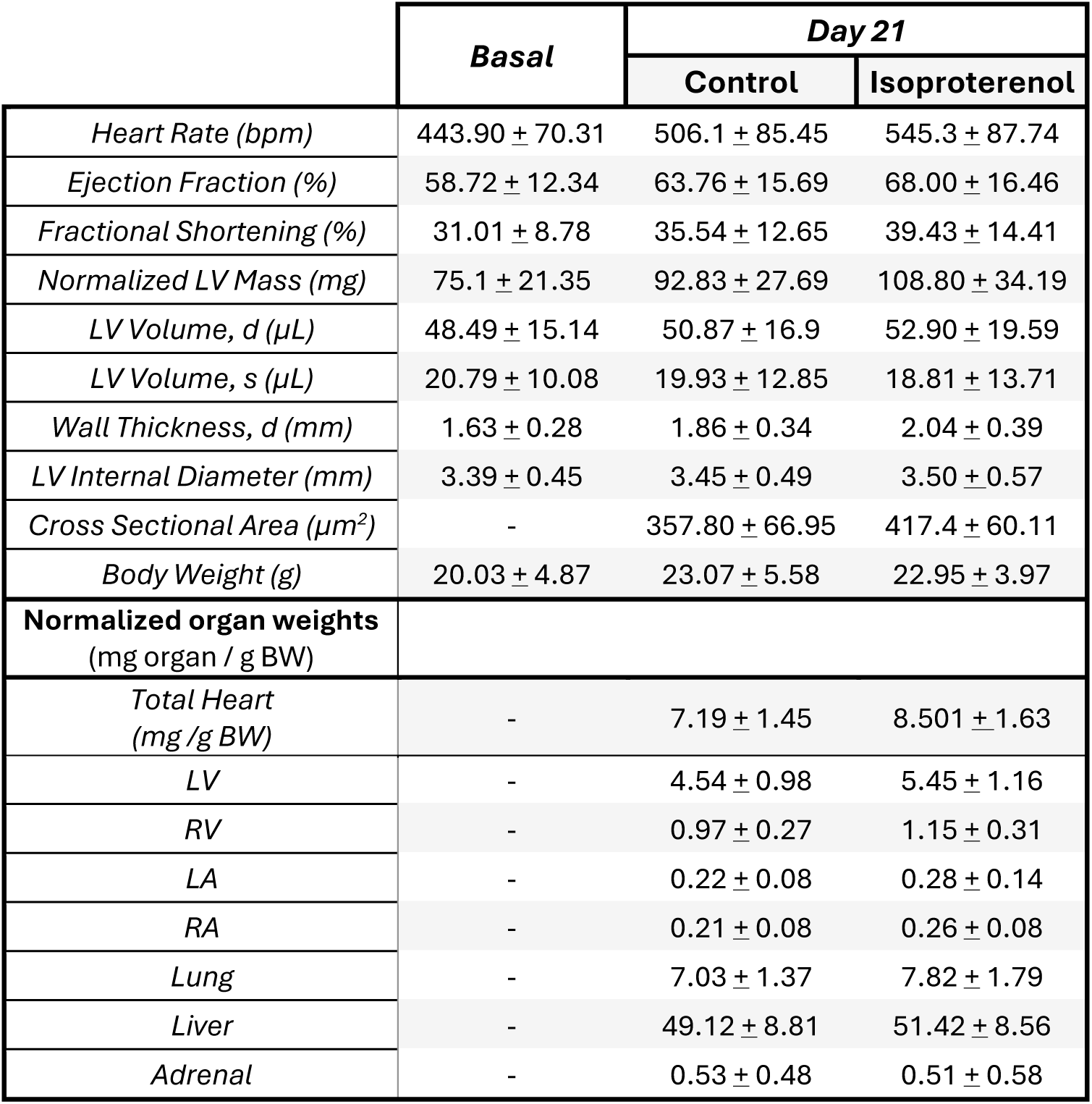
Summary of observed phenotypes in CC control and treated groups. LV = Left Ventricle, RV = Right Ventricle, LA = Left Atrium, RA = Right Atrium, BW = Body Weight, d = diastole, s = systole

|  | **Basal** | **Day 21** |  |
| --- | --- | --- | --- |
|  |  | **Control** | **Isoproterenol** |
| *Heart Rate (bpm)* | 443.90 ± 70.31 | 506.1 ± 85.45 | 545.3 ± 87.74 |
| *Ejection Fraction (%)* | 58.72 ± 12.34 | 63.76 ± 15.69 | 68.00 ± 16.46 |
| *Fractional Shortening (%)* | 31.01 ± 8.78 | 35.54 ± 12.65 | 39.43 ± 14.41 |
| *Normalized LV Mass (mg)* | 75.1 ± 21.35 | 92.83 ± 27.69 | 108.80 ± 34.19 |
| *LV Volume, d (μL)* | 48.49 ± 15.14 | 50.87 ± 16.9 | 52.90 ± 19.59 |
| *LV Volume, s (μL)* | 20.79 ± 10.08 | 19.93 ± 12.85 | 18.81 ± 13.71 |
| *Wall Thickness, d (mm)* | 1.63 ± 0.28 | 1.86 ± 0.34 | 2.04 ± 0.39 |
| *LV Internal Diameter (mm)* | 3.39 ± 0.45 | 3.45 ± 0.49 | 3.50 ± 0.57 |
| *Cross Sectional Area (μm<sup>2</sup>)* | - | 357.80 ± 66.95 | 417.4 ± 60.11 |
| *Body Weight (g)* | 20.03 ± 4.87 | 23.07 ± 5.58 | 22.95 ± 3.97 |
| **Normalized organ weights**<br>(mg organ / g BW) |  |  |  |
| *Total Heart*<br>(mg /g BW) | - | 7.19 ± 1.45 | 8.501 ± 1.63 |
| *LV* | - | 4.54 ± 0.98 | 5.45 ± 1.16 |
| *RV* | - | 0.97 ± 0.27 | 1.15 ± 0.31 |
| *LA* | - | 0.22 ± 0.08 | 0.28 ± 0.14 |
| *RA* | - | 0.21 ± 0.08 | 0.26 ± 0.08 |
| *Lung* | - | 7.03 ± 1.37 | 7.82 ± 1.79 |
| *Liver* | - | 49.12 ± 8.81 | 51.42 ± 8.56 |
| *Adrenal* | - | 0.53 ± 0.48 | 0.51 ± 0.58 |

A hallmark driver of cardiac hypertrophy and heart disease progression is the hyperactivation of the β-adrenergic signaling cascade^41^. To mimic this common HF pathway, we treated mice with ISO, a nonselective β-adrenergic agonist, to produce heterogenous cardiac stress responses based on the strains’ underlying genetic differences. Mice from each strain of the CC were divided into control (one male and one female) and treatment groups (2 male and 2 female). The treatment mice were implanted with an Alzet micropump and subjected to 30 mg/kg body weight/day of ISO, with control mice receiving pumps containing saline. After 3 weeks, ECHOs were performed again and cardiac tissue harvested for physiological weights, histology, and RNA isolation for downstream RNA-seq analyses (**Figure 1B**). To assess ISO induced changes to cardiac function, we analyzed the 21-day ECHO results in comparison to our baseline ECHO observations. ISO induced a range of phenotypic outcomes (**Figure 1D**), with most strains demonstrating a hypertrophic state as measured by increased ejection fraction, although some strains (notably CC18, CC45, and CC43) showed progression towards heart failure by markedly decreased ejection fraction. ISO-induced changes in wall thickness ranged from a decrease in some strains (e.g. A/J - 0.117 mm), suggesting the development of a dilated cardiomyopathy phenotype, to a large increase in others (e.g. NZO/HILtJ 0.881 mm), representing a hypertrophic cardiomyopathy phenotype (**Figure 1E**). All weighed tissues, on average, showed an ISO-induced increase in size when compared to control, suggesting that impaired cardiac function was leading to edema of the liver and lungs. One notable exception to this trend was the adrenal gland, which, on average, shrank slightly, consistent with prior observations that ISO stimulation results in smaller adrenal glands in some mouse strains due to attempts to reduce the unexpected (and, at least initially, unnecessary) increase in catecholamine levels in the blood^21^. Total heart weight to body weight ratio (hypertrophy) and lung weight to body weight ratio (a measurement of edema) in particular demonstrated strain-specific responses (**Figure 1F** and **G**). Ultimately, all tissue weights showed significant variation due to the interaction of drug and strain background, while sex effects were observed for total heart (females 4% higher, p = 0.02), lung (females 13% higher, p = 2.0⨯10^-9^), liver (females 4% higher, p = 6.14⨯10^-50^) and adrenal weight (females 22% higher p = 0.04) when normalized to body weight. Histological assessment of cardiomyocyte area by wheat germ agglutinin staining (**Figure 1H**) confirmed the hypertrophic phenotype observed in the whole heart weights, which significantly varied by strain and treatment status. Together, these results suggest that inherited elements play a role in the variation of cellular, tissue, and organism-level responses to ISO treatment in the CC.

### Genome Scans of Heart-Failure-Phenotypes

To connect the genetic backgrounds of our cohort to their observed phenotypes, we performed quantitative trait loci (QTL) mapping. This approach modeled each normalized trait as the sum of effects of biological sex and haplotype at each genomic position using the miQTL R package^34,35^. After setting genome-wide significance thresholds for each trait with permutation testing, we found 32 traits associated with various CC haplotypes. **Figure 2A** shows a representative genome scan for measurements of total heart weight in the control group, showing a singular significant locus on chromosome 2. In total, we observe 49 distinct genome-wide significant loci for these traits, spread across 13 chromosomes. Most of these loci (38/49, 78%) overlap with at least one other locus from a different phenotype, suggesting that common genetic drivers may coregulate many of our observed traits. We next consolidated these overlapping loci, leaving us with 9 multi-trait and 11 single-trait loci (**Figure 2B**, **Table 2**, **Supp. Table 7**).

**Fig 2.**
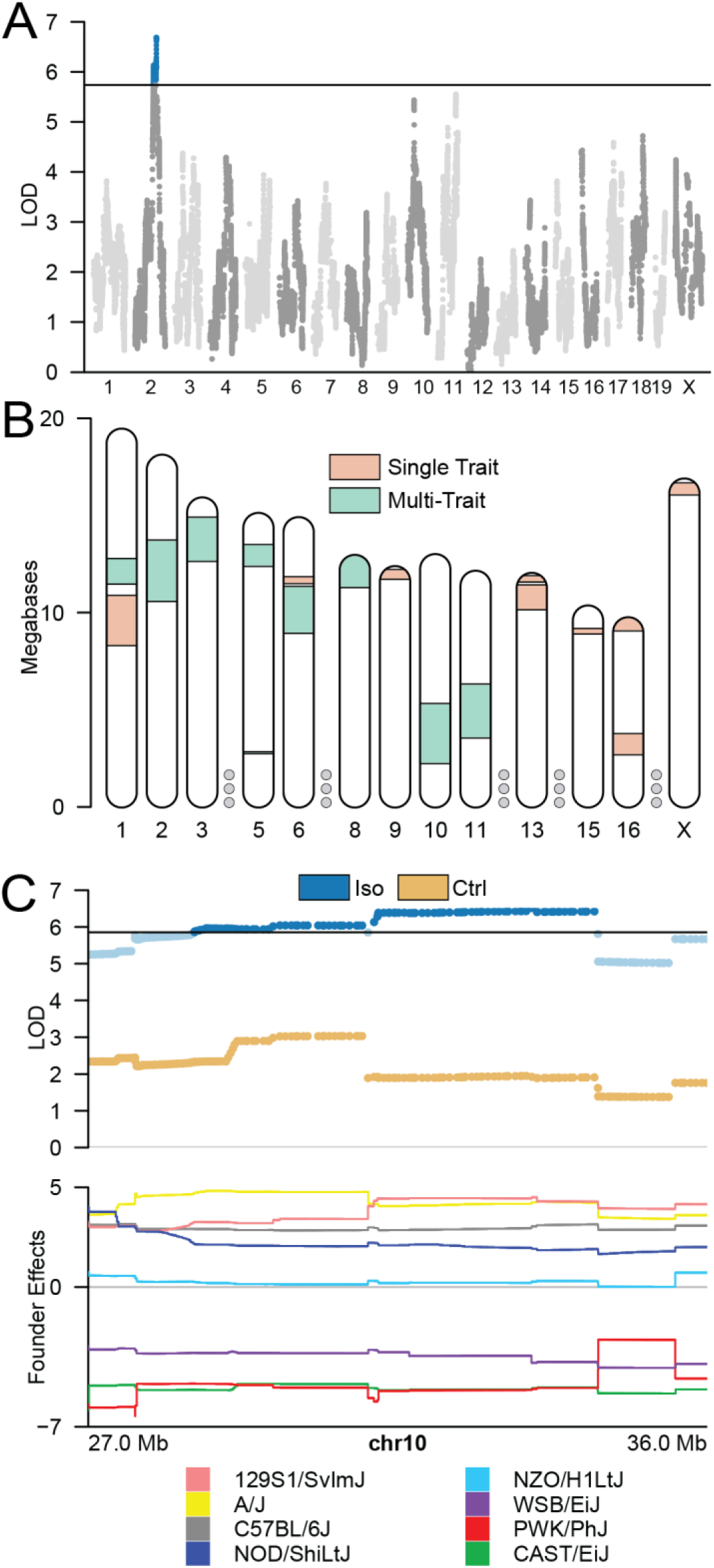
Quantitative trait locus mapping of heart failure traits in the CC. a) Genome-wide scan results for total heart weight in control group after vehicle treatment. Horizontal black line shows the threshold for significance by permutation testing, with signals exceeding that colored in blue. LOD = logarithm of odds. b) Ideogram of all loci detected in QTL mapping. Loci with overlapping significance boundaries were consolidated into 9 multi-trait regions (green) compared to 11 single-trait loci (pink). Chromosomes with no significant loci indicated by grey dots between other chromosomes c) LocusZoom plot showing association of genomic markers with left ventricular volume (top panel) in ISO-treated mice (non-significant = light blue, significant = dark blue) and control (yellow) as part of the multi-trait locus on Chr10. (Bottom panel) Inferred contributions from the eight CC founder strains show diverging directions of effect between the CAST/EiJ (green), PWK/PhJ (red), and WSB/EiJ (purple) and the remaining five founders.

**Table 2:**
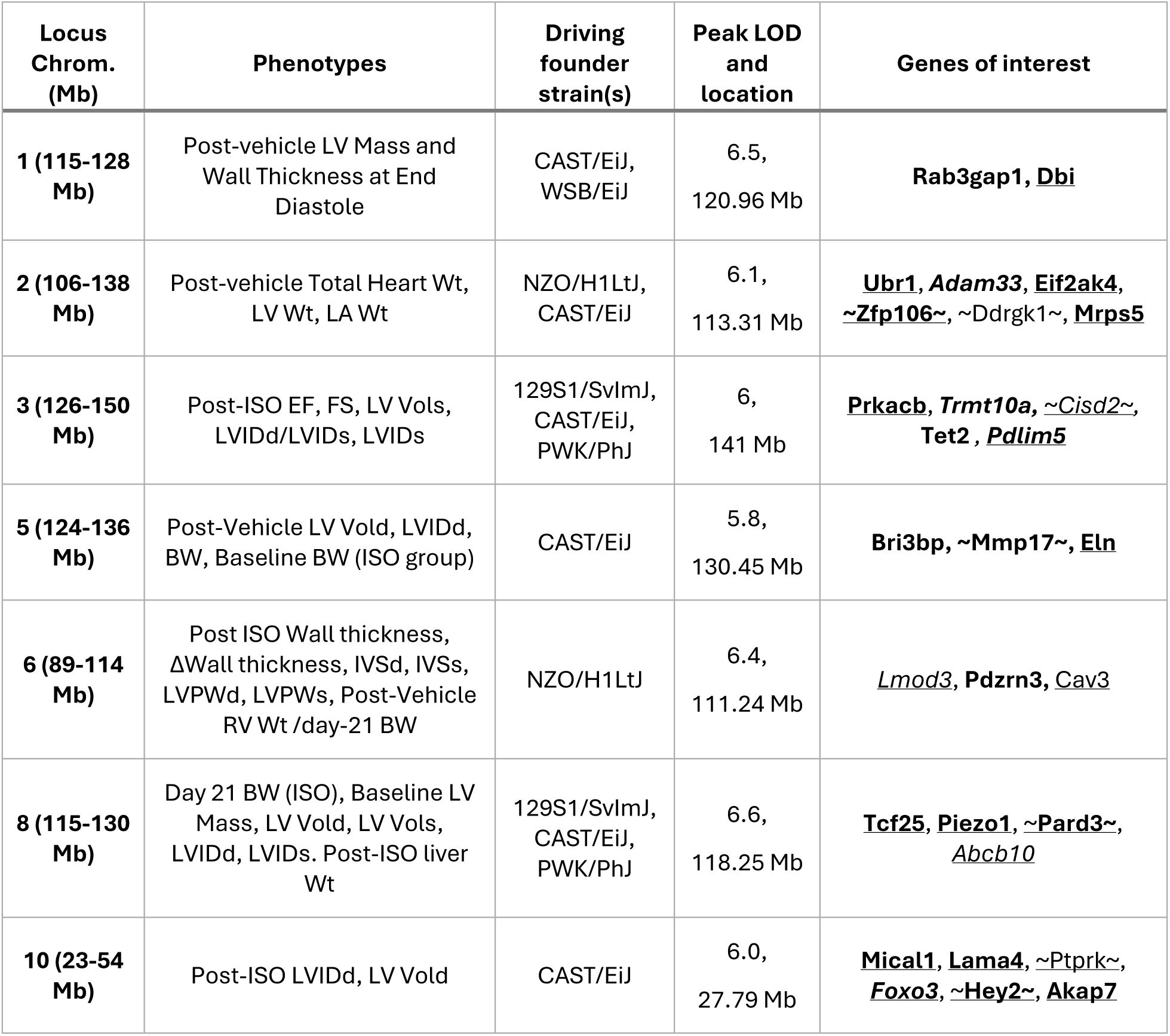
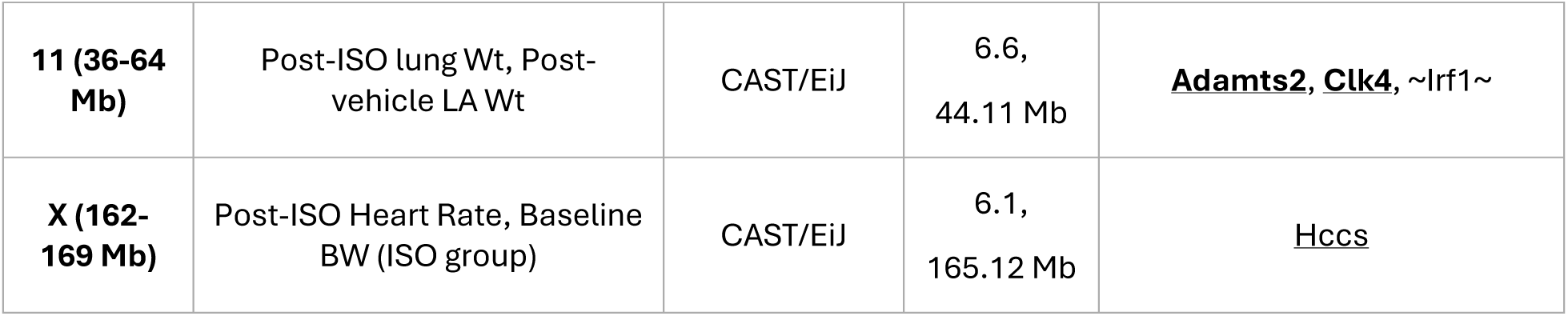
List of significant multi-trait loci along with in-locus promising candidate genes. **Abbreviations of cardiac phenotypes**: LV = Left Ventricle, LA = Left Atrium, Wt = weight, BW = Body Weight, EF = Ejection Fraction, FS = Fractional Shortening, Vol = Volume, d = Diastole, s = Systole, LVID = Left Ventricular Internal Diameter, IVS = Interventricular Septum thickness, LVPW = left ventricular posterior wall thickness. **Gene of interest annotation:**
- **Bold** denotes relevant non-synonymous SNP in founder line(s) that drive the phenotype.
- *Italic* denotes significant differential expression with ISO treatment.
- ∼ ∼ denotes presence of a cis-eQTL
- <u>Underline</u> denotes reported evidence of cardiovascular phenotype in existing knockout models or functionally relevant in muscles
- 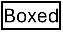 denotes well-known genes in HF context

| Locus Chrom. (Mb) | Phenotypes | Driving founder strain(s) | Peak LOD and location | Genes of interest |
| --- | --- | --- | --- | --- |
| **1 (115-128 Mb)** | Post-vehicle LV Mass and Wall Thickness at End Diastole | CAST/EiJ, WSB/EiJ | 6.5, 120.96 Mb | **Rab3gap1**, **Dbi** |
| **2 (106-138 Mb)** | Post-vehicle Total Heart Wt, LV Wt, LA Wt | NZO/H1LtJ, CAST/EiJ | 6.1, 113.31 Mb | **Ubr1**, **Adam33**, **Eif2ak4**, **~Zfp106~**, **~Ddrgk1~**, **Mrps5** |
| **3 (126-150 Mb)** | Post-ISO EF, FS, LV Vols, LVIDd/LVIDs, LVIDs | 129S1/SvImJ, CAST/EiJ, PWK/PhJ | 6, 141 Mb | **Prkacb**, **Trmt10a**, **~Cisd2~**, **Tet2**, **Pdlim5** |
| **5 (124-136 Mb)** | Post-Vehicle LV Vold, LVIDd, BW, Baseline BW (ISO group) | CAST/EiJ | 5.8, 130.45 Mb | **Bri3bp**, **~Mmp17~**, **Eln** |
| **6 (89-114 Mb)** | Post ISO Wall thickness, ΔWall thickness, IVSd, IVSs, LVPWd, LVPWs, Post-Vehicle RV Wt /day-21 BW | NZO/H1LtJ | 6.4, 111.24 Mb | **Lmod3**, **Pdzrn3**, **Cav3** |
| **8 (115-130 Mb)** | Day 21 BW (ISO), Baseline LV Mass, LV Vold, LV Vols, LVIDd, LVIDs. Post-ISO liver Wt | 129S1/SvImJ, CAST/EiJ, PWK/PhJ | 6.6, 118.25 Mb | **Tcf25**, **Piezo1**, **~Pard3~**, **Abcb10** |
| **10 (23-54 Mb)** | Post-ISO LVIDd, LV Vold | CAST/EiJ | 6.0, 27.79 Mb | **Mical1**, **Lama4**, **~Ptprk~**, **Foxo3**, **~Hey2~**, **Akap7** |
| **11 (36-64 Mb)** | Post-ISO lung Wt, Post-vehicle LA Wt | CAST/EiJ | 6.6,<br>44.11 Mb | **<u>Adamts2</u>, <u>Clk4</u>, ~Irf1~** |
| **X (162-169 Mb)** | Post-ISO Heart Rate, Baseline BW (ISO group) | CAST/EiJ | 6.1,<br>165.12 Mb | <u>Hccs</u> |

Having identified several genome-wide significant loci, we next prioritized the genes within each locus for further study using our predicted haplotype effects, eQTL mapping of genes within trait loci (see **Methods**), and known sequence variants. We found that several loci contained genes already implicated in cardiac pathology, and many of those genes presented strong concordant evidence within our study. Notably, the multi-trait loci on Chr10 (post-ISO LVIDd and LV volume) (**Figure 2C**), contains *Hey2*. This transcription factor is highly expressed in the heart^42,43^ and is known to mediate myocyte hypertrophy and cardiac dysfunction^44–46^. CAST/EiJ, the CC founder strain with the strongest predicted effect at the locus, has a non-synonymous deleterious SNP in the coding sequence of *Hey2*. Further, NZO and PWK, the two other founder strains with the same predicted direction and nearest magnitude of effect as CAST/EiJ on these traits at this locus, also share the same genotype at the peak marker for this locus. These effects also appear to be transcriptionally regulated, as *Hey2* was additionally detected as a cis-eQTL at this site. These findings indicate that our study’s framework can detect established HF-relevant genes through the integration of phenotypic, genotypic and transcriptomic data from the CC, independent of prior reported evidence, providing a high-confidence scaffold to extend to the discovery of novel genetic drivers of HF responses.

### Identification of Candidate Genes in CC Loci

The CC is known to contain large haplotype blocks, often spanning several megabases and containing dozens to hundreds of genes each^35^. The 49 loci in our study span an average of 12.48 Mb (ranging from 45 Kb to 31.71 Mb) and had a total of 2,162 genes within their bounds. This abundance creates additional barriers to the gene-prioritization process compared to other population studies. For example, human fine mapping often localizes signals to <100kb^47^ and the loci in the Hybrid Mouse Diversity Panel, another murine genetic reference population, average around 2 Mb^48^. To manage this abundance of potential candidates, we built and implemented a robust filtering system to nominate genes for functional follow-up. See **Figure 3** for a visualization of our selection workflow.

**Figure 3.**
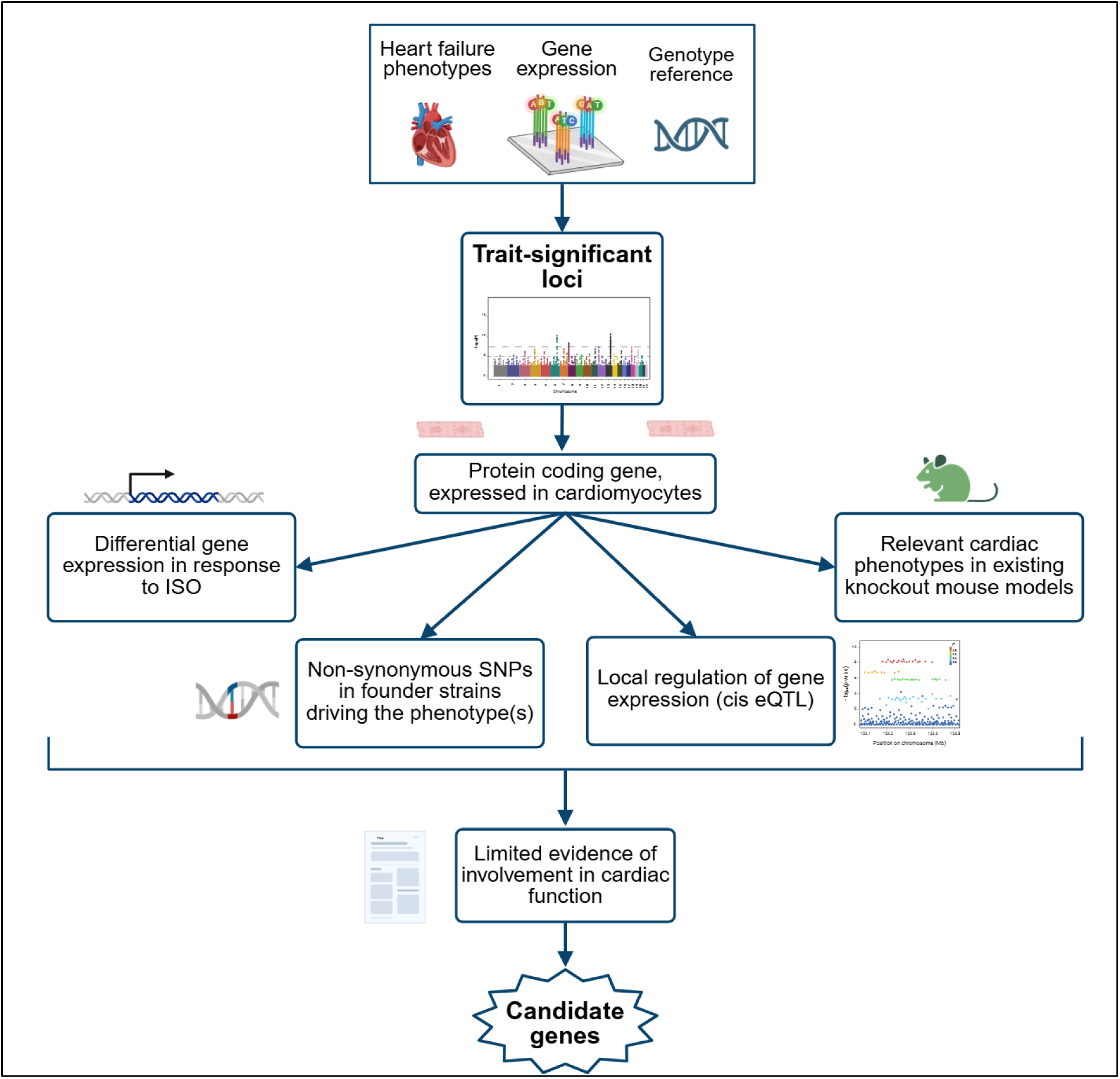
Gene candidate selection funnel. Protein-coding and NRVM-expressed genes from each significant locus were selected. Four additional key points of evidence were examined in each gene: *1)* Is the gene differentially expressed in response to ISO? *2)* Does the founder strain that drives the locus’ associated trait(s) have a non-synonymous mutation within the gene? *3)* Does the gene have a cis-eQTL within the locus? *4)* Are there knockout mouse models with reported relevant cardiovascular phenotypes? Genes with multiple points of evidence were prioritized. Well-known HF-implicated genes are then filtered out. Final candidates for downstream analyses are selected based on evidence from our data and its novelty in HF context.

We began by incorporating evidence gathered from our CC study itself. Within each trait-significant locus, we prioritized genes that demonstrated: (i) association with multiple cardiac traits in the pleiotropic loci, (ii) overlap of a significant gene cis-eQTL with the primary trait QTL, (iii) differential expression in ISO vs. control, and (iv) founder-specific coding or regulatory variation concordant with the locus’ haplotype effects (described in **Table 2**). This approach highlights any present links between genotypes, gene expression and phenotypes at a particular genome position.

Because our goal was to nominate tractable, cell-autonomous targets for downstream testing, we next restricted our candidates to protein-coding genes with sufficient expression (≥5 CPM)in our intended *in vitro* cardiomyocyte model, NRVMs. Also, we excluded genes without an obvious orthologue between mice, rats, and humans. We also prioritized genes from loci linked to traits that are more readily modeled in NRVMs (e.g. hypertrophic growth and related morphometric readouts).

To understand the relevance of the remaining candidate genes, we searched databases for reported phenotypes of mouse models for each gene. We first queried Mouse Genome Informatics (MGI)^49^ for evidence that candidate gene manipulation leads to changes in cardiovascular phenotypes (e.g. abnormal sarcomere morphology). We also searched Open Targets and surveyed the literature for any known links to cardiac or other muscle tissues in humans. In both cases, the ideal gene would have some degree of documentation regarding involvement in cardiovascular phenotypes yet not be previously implicated in heart failure or cardiac hypertrophy.

Applying this filter set yielded a shortlist of 33 genes enriched for convergent signals (see **Table 2** for a detailed list of high-confidence candidate genes at each multi-trait locus). For example, *Adamts2* at the chromosome 11 locus (atrial weight, lung edema) carries a deleterious nonsynonymous variant in the CC founder strain predicted to have the strongest effect at this locus. It also has prior evidence linking it to cardiac hypertrophy and edema from our own prior research in the HMDP^50^; we therefore considered it a high-confidence candidate. Similarly, *Manba,* at the chromosome 3 locus (post-ISO cardiac function and heart wall dimensions) shows a cis-eQTL and founder-specific nonsynonymous variation (CAST/EiJ, PWK/PhJ). Until now, the only evidence of *Manba*’s involvement in a cardiovascular trait is from a GWAS of atrial fibrillation (AGES-Reykjavik study), where it presented as a candidate causal gene^51^. Due to the sparsity of the gene’s functional information, we did not highly prioritize *Manba* for subsequent *in vitro* investigation despite it being indicated by our internal data.

### *In vitro* validation of locus hits

Based on the gene selection criteria described above, we chose ***Abcb10***, ***Mrps5*** and ***Lmod3*** as our candidates for *in vitro* validation (see **Table 3** for detailed summary of supporting evidence for these three genes). As β-adrenergic stimulation’s hallmark is cardiomyocyte hypertrophy, we used cell size from isolated neonatal rat ventricular myocytes (NRVMs) as our primary readout for the response to ISO stimulation with or without the knockdown of our gene of interest using siRNAs. ISO’s engagement of the hypertrophic pathway in our cells was confirmed at a molecular level by qPCR in non-transfected NRVMs by a 4.6 fold upregulation of *Nppb* (p=0.03), a canonical myocyte hypertrophic marker^52^ (**Supp. Figure 1**), while its physical effect on cell size was confirmed through quantification with the CellPose program (See **Fig. 4a** for representative NRVM images)

**Figure 4.**
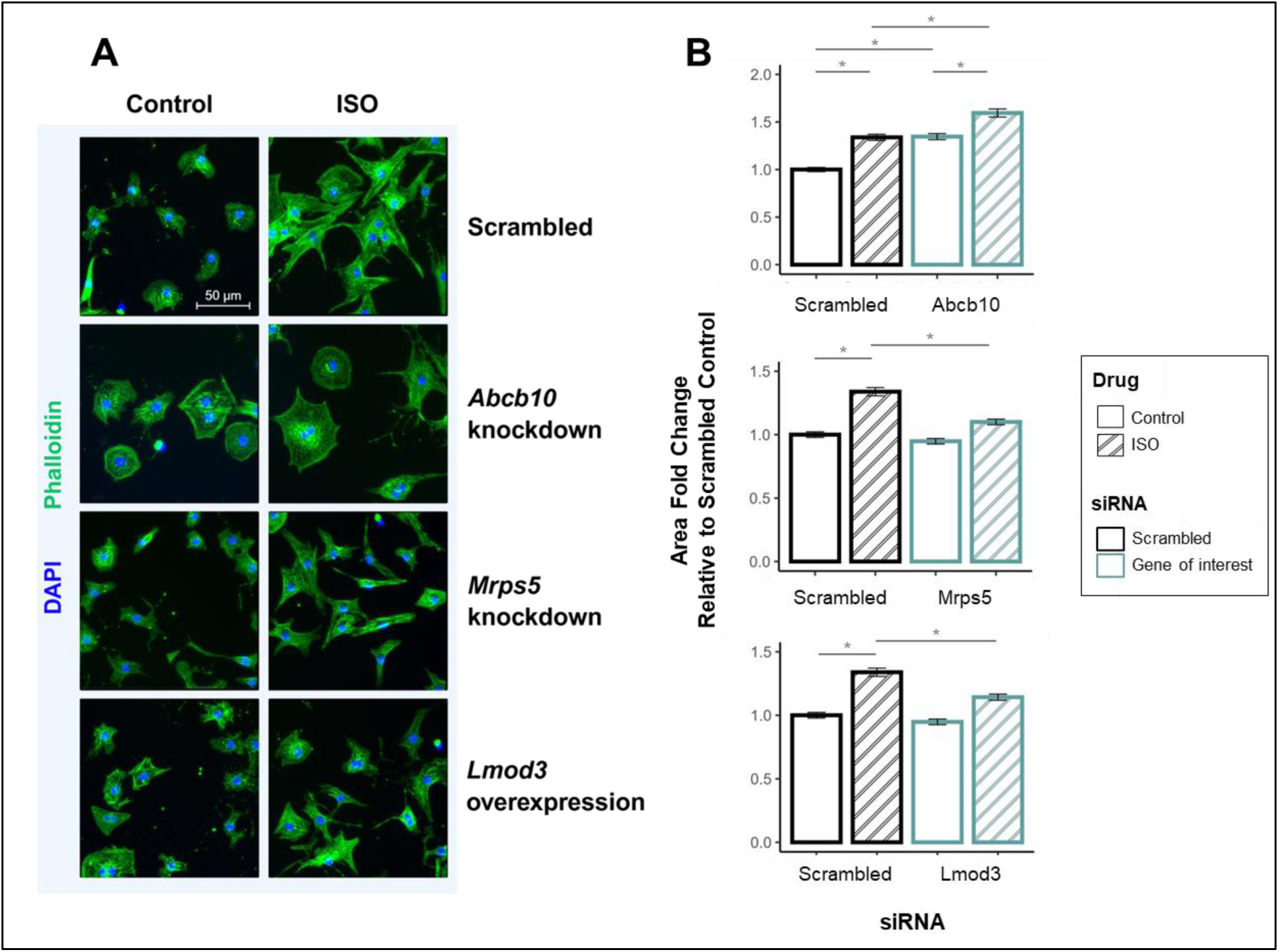
Abcb10, Mrps5 and Lmod3 are involved in the regulation of cardiomyocyte area. NRVMs were transfected with a scrambled sequence or siRNA of *Abcb10, Mrps5* or *Lmod3* and incubated for 24 hours, followed by ISO treatment in DMEM-ITS for 48 hours and subsequent cell area quantification. **A)** Representative images of NRVMs from each experimental group (gene knockdown vs ISO challenge). Cell nuclei and actin networks were visualized via DAPI and Phalloidin staining, respectively. **B)** Quantification of cell area of each group relative to the control scramble-treated group within each study. *Abcb10* knockdown increases cell size globally, *Mrps5* and *Lmod3* knockdown attenuate ISO-induced hypertrophy. Data are presented as mean ± 2*SEM (N=9 from three separate experiments with triplicates). * indicates p<10⨯10^-12^ (post hoc TukeyHSD test following 2-way ANOVA) for selected comparisons. Each experimental group has between 1300-1700 cells recorded for cell size quantification.

**Table 3.** Table of evidence for selected candidate genes.

| Gene (chromosome) | Associated Phenotypes | cis eQTL | Differential Expression | Non-synonymous SNP in driving strain(s) | Reported cardiac-relevant phenotypes in knockout model (MGI database) | CPM in NRVMs | Evidence in the literature |
| --- | --- | --- | --- | --- | --- | --- | --- |
| **Abcb10 (8)** | Baseline LV Vold, LVIDs, LV mass<br>Post-treatment liver mass (ISO-treated) | No | Yes (log2FC 0.15, p 0.0026) | No | Increased response of heart to induced stress<br>abnormal mitochondrial physiology | 20.38 | Heterozygous functional mouse (Abcb10<sup>+/-</sup>) have impaired systolic and diastolic function and reduced recovery subsequent to ischemia/reperfusion injury<sup>45</sup><br>Cardiomyocyte-specific deletion causes increased ROS production, mitochondrial structural abnormalities and cardiac fibrosis<sup>58</sup> |
| **Lmod3 (6)** | ISO-treated IVSs, LVPWd, LVPWs, total wall thickness | No | Yes (log2 fold change 0.15, p=0.01) | No | Decreased cardiac muscle contractility<br>Abnormal sarcomere morphology | 5 | Dysfunctional protein causes thin filament disorganization and nemaline myopathy<sup>63</sup> |
| **Mrps5 (2)** | LA weight, LV weight, total heart weight (control-treated) | No | No | CAST |  | 42 | Single allele deletion model showed increase in CM proliferation and cardiac regeneration post-MI<sup>61</sup><br>Constitutive knockout causes embryonic lethality; postnatal knockout induces cardiac hypertrophy and heart failure<sup>60,61</sup> |

The heart is the most mitochondria-dense tissue due to its high-energy demand and constant contractile activity as a muscle^53^. On a chromosome 8 locus associated with baseline heart wall dimension, baseline ventricular function and post-ISO edema indicators^21,54,55^ (see **Table 2**), we identified *Abcb10,* a gene encoding an ATP-binding cassette (ABC) transporter on the mitochondrial membrane as the most likely candidate gene at this locus. *Abcb10* was significantly upregulated with ISO (log2 fold change of 0.15, p=0.0026, although strain-to-strain responses varied, such as CC015 increasing by 2.2 fold and CAST/EiJ decreasing by half, suggesting a strain-specific upregulation of this gene in response to ISO stimulation). *Abcb10* is known to regulate oxidative stress response^56,57^, as *Abcb10* deficiency in rodents detrimentally affects mitochondrial function^58^, and impairs post-ischemia/reperfusion injury recovery^45^. In our study, without siRNA-mediated knockdown (scramble siRNA), we observe an average increase of NRVM size of 34% in response to ISO (p=3.5⨯10^-12^). Compared to its control scramble-treated group, a knockdown of *Abcb10* expression by 51% (p=0.013, **Supp. Figure 2**) increases NRVM size by 35% (p=3.5⨯10^-12^) without ISO stimulation and even more so with ISO by 59% (p=3.5⨯10^-12^). **(Figure 4).** This suggests that *Abcb10* knockdown mimics the hypertrophic effects of ISO (P=0.98 between scramble ISO and *Abcb10* Control), but further stimulation of the β-adrenergic signaling cascade remains possible after ISO treatment (an additional 19% increase, p=3.6⨯10^-12^)**(Figure 4, Supp. Figure 3)**.

Another mitochondrial-related genes we have shortlisted is *Mrps5*, encoding mitochondrial ribosomal protein S5, a known regulator of mitochondrial respiration^59^. *Mrps5* resides within a chromosome 2 locus that is significantly associated with control total heart and chamber weights, with CAST/EiJ as the phenotype-driving founder strain (see **Table 2**). To date, studies on *Mrps5*’s role in the heart remain sparse, with only one study finding *Mrps5* critical for embryonic heart development and postnatal cardiac function, while a separate group demonstrated that *Mrps5* reduction can mediate post-MI cardiac regeneration^60,61^. In our study, we have successfully knocked down *Mrps5* expression by 85.6% **(Supp. Figure 2)**. This significant decrease in *Mrps5* expression mitigates the ISO-induced hypertrophic response. Compared to the control scramble-treated group, ISO increases cell size by 34% (p=2.6⨯10^-12^), while ISO along with decreased *Mrps5* only increases cell size by 10% (p=3.57⨯10^-8^) **(Figure 4)**. Between the *Mrps5* knockdown groups, ISO increases cell size by 15% relative to control, half of what was observed in the scramble-treated NRVMs (p=1.22⨯10^-13^)**(Supp. Figure 3)**.

In a muscle tissue such as the heart, actin dynamics are essential to organ function and can be disturbed by excessive stress signals such as chronic β-agonism, leading to altered myocyte morphology and contractility^62^. A chromosome 6 locus was significantly associated with post-treatment ventricular weight and heart wall dimensions (see **Table 2**). Within this locus resides *Lmod3*, encoding Leiomodin-3 protein, which canonically regulates actin filament formation and sarcomere integrity^63,64^. We suspect *Lmod3* may help regulate cardiac muscle growth in the context of β-adrenergic stimulation. Paradoxically, transfection with *Lmod3*-targeting siRNAs increased *Lmod3* expression by 140% (p=0.038). We validated this upregulation by Western blot as well, showing a 67% increase (p=0.022) in protein abundance after siRNA administration (**Supp. Figure 2**). As previously reported for other genes^65,66^, this unexpected phenomenon might be due to compensatory gene upregulation following transient gene knockdown, or unintended siRNA-promoter binding (also known as “RNA activation”). Regardless of the cause, our results show that increased *Lmod3* expression partially attenuates ISO-induced hypertrophy. Compared to the control scramble -treated group, ISO increases cell size by 34% (p=1.2⨯10^-13^), while ISO in the presence of elevated *Lmod3* only increases cell size by 14% (p=1.88⨯10^-13^) **(Figure 4)**. Since *Lmod3* is downregulated in ISO-treated NRVMs, we suspect our inadvertent *Lmod3* over-expression has strengthened *Lmod3*’s inhibitory function in the β-adrenergic hypertrophic pathway in NRVMs (**Supp. Figure 1**).

## Discussion

The goal of our study was to leverage the diversity of the Collaborative Cross to identify novel genetic loci associated with cardiac physiology and pathophysiology through variation in β-adrenergic stress responses. The larger genetic variability of the CC, compared to other Genetic Reference Populations (GRPs), results in a wider distribution of cardiac phenotypes at baseline and in more significant variation in their response to catecholamine challenge compared to prior GRP HF studies (see, for example, heart weight to body weight ratio in the CC vs the HMDP^21^ in **Supp. Figure 4**). In this study, we identified 49 genomic loci associated with heart-failure-relevant phenotypes, with 78% of these loci associated with multiple traits. This highlights that the complexity of cardiac phenotypic interplay may be driven by distinct genomic loci.

A challenge of relating QTL signals to candidate genes in animal GRPs is that the resolution of these panels are typically significantly larger than that seen in human cohorts. Where human populations often have resolutions for loci that are in the 500kb or smaller range^47^, many GRPs measure their loci in terms of megabases^48,67,68^. Indeed, we observe large loci in the CC, with an average locus size of 12.83 Mb, consistent with past CC studies^12,35,36^. These large locus sizes are one of the reasons for the development of the Diversity Outbred (DO) population, the product of random breeding of mice derived from the same founder lines of the CC and now maintained in a highly heterogenous outbred stock. Although the DO has sub-Mb resolution^69^, its construction means that its genomes are not fixed–each DO mouse is unique and must be individually genotyped and analyzed. One consequence of this is that our experimental design, in which genetically identical mice were subjected to either ISO or saline, is not possible in the DO.

Given the large loci observed in our study, it was imperative that we use a robust filtering approach (**Figure 3**) to narrow down the candidate genes to a minimal set of high-confidence candidates. Our filtering framework incorporated existing genotyping information from the CC, *in vitro* NRVM data, study-specific transcriptomic results, and the phenotype databases of MGI and Open Targets. This allowed us to confidently sort through the dozens-to-hundreds of genes within each locus and narrow them down to a small handful of testable candidates. Using this framework, we identified multiple genes which we linked to cardiac hypertrophy and HF progression. These genes were comprised of both well studied and novel candidates (see **Table 2**), spanning a variety of cellular functions such as oxidative stress (*Mrps5, Abcb10*), actin dynamics (*Lmod3*), cell migration (*Pdlim5*), calcium handling (*Pde4*), collagen processing (*Adamts2*), transcriptional regulation (*Hey2*) and more. We have confirmed that three genes (*Mrps5, Abcb10, Lmod3*) which were previously understudied in HF, regulate cardiomyocyte phenotypes using *in vitro* models, demonstrating in NRVMs that manipulating the expression levels of these candidate genes alters myocyte hypertrophic response to ISO and/or cell dimensions at baseline. This demonstrates that these genes participate, either directly or indirectly, in the β-adrenergic signaling cascade and in cardiac hypertrophy modulation.

ISO canonically activates cyclic-AMP-protein kinase A (cAMP-PKA) signaling, increases intracellular calcium and subsequently acutely enhances cardiomyocyte contractility and, after chronic stimulation, induces cell morphological remodeling^70^. One driving feature of this remodeling is sarcomeric organization. *Lmod3* mutations can cause generalized skeletal muscle weakness^63,64,71^, but the role of *Lmod3* in cardiac muscle remains unexplored despite its high expression in the heart^71^. As *Lmod3* regulates actin nucleation and filament formation^63,64^, overexpressing *Lmod3* in an ISO-stressed environment may partially rescue actin destabilization and dampen some ISO-driven remodeling effects in cardiac muscle cells.

ISO can also induce mitochondrial dysfunction, calcium mishandling and oxidative stress, leading to activation of various mitogen-activated protein kinases (MAPK) which contribute to pathologic hypertrophy^72–75^. The hypertrophic effect we observed when silencing *Abcb10* in NRVMs may be explained by its known role in protecting against oxidative stress. One group showed heterozygous mice (Abcb10^+/-^) had impaired systolic and diastolic function and reduced recovery from ischemia/reperfusion injury when compared to WT. Another group showed that cardiomyocyte-specific *Abcb10* knockouts without any pathological stimulation exhibited increased cardiac fibrosis and mitochondrial dysfunction, likely via a ferroptosis-dependent mechanism^58^. It is possible that *Abcb10*-deficient NRVMs in our study experienced heightened cellular stress by accumulation of reactive oxygen species, mimicking the pro-hypertrophic effect of ISO stimulation.

In contrast, *in vivo* reduction of *Mrps5* promotes cardiomyocyte proliferation and heart generation via compensatory upregulation of activating transcription factor 4 (ATF4)^60,61^. ATF4 is activated by multiple cardiac stress signals and serves as an cardioprotective oxidative stress inhibitor^76^. This may explain why our data show blunting *Mrps5* expression can partially dampens ISO’s pro-hypertrophic effect, although the role of Mrps5 in delicately balancing physiological and pathologic hypertrophy is still unclear.

Our study has several limitations. We were resource-limited to 6 mice per strain (2 control, 4 treated) This lowered our sensitivity to intra-strain variability caused by experimental noise and any latent heterozygosity which may still exist within some of the CC lines^77^. The addition of more mice per strain could constrain this variability. Despite this limitation, it is important to note that genome scans do not operate on a strain-by-strain basis when identifying associations, but on a founder haplotype by founder haplotype basis. That is, if we assume that every locus contains equal contributions from each founder strain, we have approximately 8-9 strains, or an N of 48-54 animals with each haplotype at each site. If two founder lines share ancestry at a specific locus, we may have an even larger effective sample size for that specific locus. Another limitation is the limited number of strains in the CC population, which was originally envisioned to be much larger before running into significant bottlenecks during inbreeding^78^, which results in larger LD blocks and limited resolution. Despite these limitations, we were able to recover a significant number of loci and identify candidate genes within them. Future efforts could focus on adding additional strains, such as mice from other inbred GRPs, or supplementing the CC data with mice from the Diversity Outbred population to increase power and improve mapping resolution. Finally, we acknowledge that, although a commonly used and well-established *in vitro* model for cardiac hypertrophy, NRVM monoculture is an imperfect model for the 3D heart and, furthermore, cannot identify contributions from other cardiac cell types which may be expressing the genes of interest. Larger, more comprehensive studies are needed to fully validate and understand the mechanisms that underlie the candidate genes we identified within our loci.

In conclusion, our study expands on prior work done in the CC to explore how diverse mice respond to a common hypertrophy-inducing stressor and identifies novel genetic variations contributing to cardiac disease. *Mrps5, Lmod3* and *Abcb10*, along with the other candidate genes selected by our filtering approach presents promising avenues for further mechanistic dissection of common, pleiotropic pathways in cardiac dysfunction. Our data represents a resource for other researchers in terms of candidate genes and animal models for investigating heart disease. Uncovering novel genes without established association with heart disease suggest possible novel targets for pharmacological and therapeutic interventions and/or additional factors to assess heart disease risk propensity in humans.

## Data and Code Availability

Phenotype data is available through Mendeley Data at 10.17632/vyf5x4ygrv.1. Bulk RNAseq data for NRVMs is available at the Sequence Read Archive (BioProject ID: 1328793). Bulk RNAseq data for CC mice is available at the Sequence Read Archive (BioProject ID: 1331689). Code is available at github.com/RauLabUNC/cc_gwas

## Author Contributions (CRediT)

Conceptualization – CDR

Data Curation – THK, ANL, BG, CDR

Formal Analysis – THK, ANL, BG, AH

Funding Acquisition - CDR

Investigation – THK, ANL, BG, CL, AH, SR, AA, AD, SA, ELS, LGK, MG, CDR

Methodology – THK, ANL, BG, CDR

Project Administration – BCJ, RBB, CDR

Resources – THK, ANL, BG, CL, CDR

Software – ANL, BG, AH, AD, CDR

Supervision – BCJ, RBB, CDR

Validation – THK, ANL, BG, AH

Visualization – THK, ANL, BG

Writing: Original Draft – THK, ANL, BG, CDR

Writing: Review & Editing- All Authors

## Acknowledgements

We used BioRender in figures 3 and 4.

